# Mechanistic Evaluation of Amplification Lag in Paper-Based Colorimetric Loop Mediated Isothermal Amplification (LAMP) and Its Reduction by BSA Pre-Coating

**DOI:** 10.1101/2025.10.25.684418

**Authors:** Taehong Kim, Gopal Palla, Bibek Raut, Mohit S. Verma, Arezoo M. Ardekani

## Abstract

Colorimetric loop-mediated isothermal amplification (LAMP) on microfluidic paper-based analytical devices (μPADs) offers a low-cost, disposable, and equipment-free alternative to liquid LAMP assays. However, amplification on μPADs is consistently slower, by 5–46%, than reactions in tubes. To identify the origin of this delay, we evaluated heat transfer, diffusion in porous cellulose, and nonspecific adsorption of LAMP components across both high- and low-copy input regimes. Our results show that once thermal equilibrium is reached, reduced effective diffusion is the dominant contributor to the kinetic lag at low copy numbers, whereas nonspecific adsorption becomes the primary barrier at higher template concentrations. Pre-coating the paper with bovine serum albumin (BSA) mitigates adsorption. It narrows the tube-to-paper gap, thereby accelerating amplification of the SARS–CoV-2 ORF7ab synthetic gene by an average of 6 minutes, from 1E3 to 1E5 copies per reaction. These findings provide a mechanistic basis for the copy-number-dependent behavior of μPAD LAMP and offer simple, low-cost strategies to improve the speed and reliability of μPAD nucleic acid assays.

## 1 Introduction

Colorimetric loop-mediated isothermal amplification (LAMP) on microfluidic paper-based analytical devices (µPADs) offers a low-cost, disposable, and equipment-free alternative to liquid LAMP assays. However, amplification on µPADs is consistently slower, by 5–46%, than reactions in tubes. To identify the origin of this delay, we evaluated heat transfer, diffusion in porous cellulose, and nonspecific adsorption of LAMP components across both high- and low-copy input regimes. Our results show that once thermal equilibrium is reached, reduced effective diffusion is the dominant contributor to the kinetic lag at low copy numbers, whereas nonspecific adsorption becomes the primary barrier at higher template concentrations. Pre-coating the paper with bovine serum albumin (BSA) mitigates adsorption. It narrows the tube-to-paper gap, thereby accelerating amplification of the SARS-CoV-2 ORF7ab synthetic gene by an average of 6 minutes, from 1E3 to 1E5 copies per reaction. These findings provide a mechanistic basis for the copy-number-dependent behavior of µPAD LAMP and offer simple, low-cost strategies to improve the speed and reliability of µPAD nucleic acid assays.

During and after the COVID-19 pandemic, nucleic acid amplification technologies (NAATs), particularly Loop-mediated Isothermal Amplification (LAMP) ^1–3^, have drawn significant attention across clinical diagnostics, environmental monitoring, and onsite biosurveillance. LAMP is a powerful isothermal amplification method that achieves high sensitivity and specificity without thermal cycling, enabling robust amplification with a strand-displacing polymerase at a single temperature ^4–6^. By employing a set of loop-forming primers that target multiple regions on the template, LAMP accelerates extension while suppressing off-target binding, and in doing so, eliminates the repeated denaturation/annealing steps required by PCR-based methods ^4,5,7^. In addition, colorimetric indicators enable direct visual readout. This colorimetric signal reduces the need for complex instruments, allowing simple heaters to suffice. However, for more reliable results, studies have included on-board image capture and image analysis to improve the sensitivity and allow quantitative interpretation of results similar to those seen with real-time quantitative polymerase chain reaction (qPCR) but without the need for fluorescence dye or optics ^8–17^. These attributes collectively position LAMP as an attractive platform technology for point-of-need molecular tests that require speed, portability, and operational simplicity.

Given these advantages, LAMP has been frequently implemented on paper-based analytical devices (µPADs) to create low-cost, field-deployable diagnostics ^2,3,9^. µPADs and paper-polymer hybrid LAMP devices integrate biochemical amplification with capillary-driven microfluidic channels, taking advantage of paper’s wicking behavior and ease of patterning ^18,19^. It is highly compatible with nucleic acids and proteins and operates with small reaction volumes. As a result, µPAD LAMP has been used to detect diverse targets, including Zika virus ^20,21^, bacterial DNA ^22^, foodborne pathogens ^23–26^, and rotavirus A ^27^. Nevertheless, translating isothermal amplification from bulk liquid to a porous fibrous matrix introduces new physical constraints, commonly manifested as non-uniform color development, delayed time-to-positive signal, and increased variability across devices ^2,9,28^.

Although LAMP kinetics and mechanisms have been examined ^29^, the reaction dynamics described in the paper remain insufficiently characterized, despite their practical relevance. Here, we compare µPAD and Tube LAMP under matched primer and reagent conditions to identify the dominant contributors to the observed kinetic lag on paper. We focus on three coupled factors: (i) heat-transfer behavior, (ii) effective diffusion in the porous matrix, and (iii) nonspecific adsorption of primers, polymerase, or template to cellulose fibers.

To standardize the substrate and facilitate reproducibility, we selected cellulose chromatography paper for its widespread availability and stable capillary properties ^30^. In addition, the device design and material comparison conducted by Davidson *et al*. demonstrated that Ahlstrom Grade 222 exhibits improved wicking behavior, more uniform reagent distribution, and a smaller reaction area, owing to its greater thickness compared with traditional Grade 1 chromatography paper. Their evaluation showed that Grade 222 supports consistent reagent drying, reliable colorimetric signal development, and minimized variability, making it a suitable substrate for µPAD RT-LAMP assays ^2^.

Heat transfer is the first design parameter in isothermal amplification. LAMP typically operates at 63-67°C, and temperature drifts can alter polymerase activity, primer-template hybridization kinetics, and pH-sensitive indicators ^31^. While heat transfer in porous media and microfluidic substrates has been broadly studied ^32,33^, few studies have directly isolated how heat transfer in µPADs influences the temperature field during LAMP amplification.

Beyond temperature control, reactant transport is critical to the growth phases of LAMP. In tube-based reactions, mixing is effectively homogeneous, whereas in µPAD systems, reactants primarily migrate by diffusion within a porous network. Prior studies have examined imbibition-driven convection and diffusion in cellulose and nitrocellulose for LAMP-compatible reactions ^30^. In our previous work, using the same LAMP formulation, we observed that amplification proceeded faster in tubes than in paper substrates ^2^. However, the transport mechanisms underlying this kinetic discrepancy were not systematically investigated.

A third factor is the nonspecific binding of biomolecules to cellulose. Cellulose-based µPADs present highly porous, high-surface-area fibrous networks that can physically adsorb nucleic acids and proteins through hydrogen bonding, electrostatic interactions, and hydrophobic effects, independent of any intended affinity chemistry ^34^. Such adsorption can deplete the free concentrations of primers or enzymes, thereby reducing reaction rates.

Despite the expanding use of LAMP on paper substrates, the mechanistic origins of its slowed and more variable kinetics in porous media remain poorly understood. As a result, design choices for µPAD-based amplification systems, such as substrate selection, device geometry, and thermal control, have often been guided empirically rather than informed by a clear understanding of the underlying physical and biochemical factors. To address this gap, the present study compares LAMP performance in matched Tube and µPAD. It evaluates the relative influence of heat-transfer behavior, diffusion-driven transport, and biomolecular adsorption within cellulose matrices. Through controlled experiments and direct measurements of reaction kinetics and substrate-dependent losses, we identify the key mechanisms that contribute to the kinetic lag frequently observed in µPAD LAMP.

These insights provide practical guidance for improving the reliability, speed, and consistency of µPAD molecular diagnostics and support the development of more robust point-of-need nucleic acid tests.

## 2 Experimental

### 2.1 Fabrication of the µPAD Device

Paper strips were fabricated following the previously published protocol ^2,14,15^. The paper device consists of a serial array of 3 mm × 3 mm × 1.1 mm (length (l) × width (w) × height (h)) paper pads and 2 mm × 3 mm × 0.5mm (l×w×h) polystyrene spacers (Tekra Double White opaque HIPS Polystyrene Litho Grade) ^3^. The Ahlstrom chromatography electrophoresis and blotting paper, grade 222, is used. It is cut with an electric leather-cutting machine (Xhixumm) and fixed with adhesive film (Tekra Melinex 454 polyester). Then, the strips with paper pads are encapsulated in a 1.5-mm-thick laser-cut Acrylic frame and a PCR adhesive film ^13^.

### 2.2 LAMP Reaction

Homemade master mix and primer mix are used following the previously reported protocol ^2,14,15^. The master mix is composed as shown in Table 1. Six primers targeting the ORF7ab (NC_045512.2) region of the SARS-CoV-2 genome are utilized, as shown in Table 2^2^. In Tube LAMP, 7.5 µL of template is added to 17.5 µL of reaction mix, for a total volume of 25 µL. In the µPAD, 7.5 µL of a mixed master mix-template solution is used per reaction. For the other LAMP experiment, to control the reaction pre-treatments, some chemicals are dried overnight (24 hours) at room temperature to ensure the liquid volume in the paper device does not exceed the device’s capacity. Previously reported water bath heaters (ThermiQuant™AquaStream and ThermiQuant™MegaScan) are used to incubate the LAMP reactions. As a target DNA, we use the SARS-CoV-2 ORF7ab (NC_045512.2) synthetic gene from GenScript.

**Table 1.** Composition of Master mix.

| Components | Concentration |
| --- | --- |
| KCl | 62.5 mM |
| MgSO<sub>4</sub> | 10 mM |
| dNTPs | 1.75 mM |
| Bst 2.0 | 807 mU/μL |
| Phenol red | 0.125 mM |
| dUTPs | 0.175 mM |
| Antartic Thermolabile UDG | 0.25 mU/μL |
| Tween 20 | 1.25% |
| Primer mix | 12.5% |
| Betaine | 25 mM |
| Trehalose | 18% |
| BSA | 0.626 μg/μL |

**Table 2.**
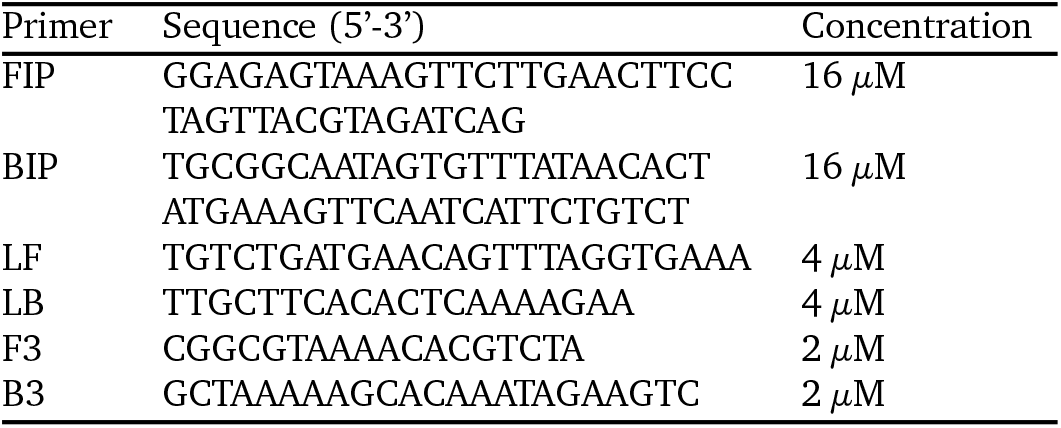
Composition of Primer mix.

### 2.3 Image Analysis and Quantitative Evaluation of Colorimetric Signals

Time-lapse images (20-30 s interval) are captured using either ThermiQuant™ MegaScan (for µPADs) or Thermi-Quant™ AquaStream (for Tubes) while incubating at 65°C in a water bath. Time-lapse images are processed using Amplimetrics™ to automatically crop the tubes or µPADs and extract hue vs time data ^13,15^.

The reaction is first classified as positive or negative using a hue threshold of 5. For positive reactions, the hue-time data are normalized to the maximum hue, and quantification time (Tq) is obtained using Amplimetrics™. Tq is defined as the time to the second derivative maxima, which correspond to the maximum reaction acceleration.

### 2.4 Temperature Measurement in Tube and µPAD

Temperatures are measured using K-type thermocouples and a multichannel data logger (AZ Instruments K-type 299 thermocouple recorder with 8G SD card, Amazon, USA). Two thermocouples are each dipped into either two tubes (a hole drilled through the PCR tube cap) containing 25 µL water or onto a wet µPAD housed in a 1.5 mm-thick acrylic frame. To prevent water from leaking into the µPAD and tubes, the thermocouple leads are glued using superglue. We start recording the temperature at room temperature, then dip the tubes and µPAD cartridges into a 65°C pre-heated water bath, and record continuously every second for 80 min.

## 3 Results

### 3.1 Comparison Between Tube and µPAD LAMP

Figure 1 illustrates the progression of the LAMP reaction by quantifying hue changes over time. Figure 1a shows tube reactions submerged in a 65°C water bath, captured 30 min after the reaction was initiated. All reactions use identical mixtures (Table 1) and contain 1.0E+03 through 1.0E+06 genomic copies per reaction. The range is chosen within the limits of quantification of µPAD based on the previous work ^15^. After 30 min of incubation, tubes containing 1.0E+06 copies exhibit a complete transition to yellow, whereas reactions with fewer copies display only partial hue changes, and NTCs retain their original color.

**Fig. 1.**
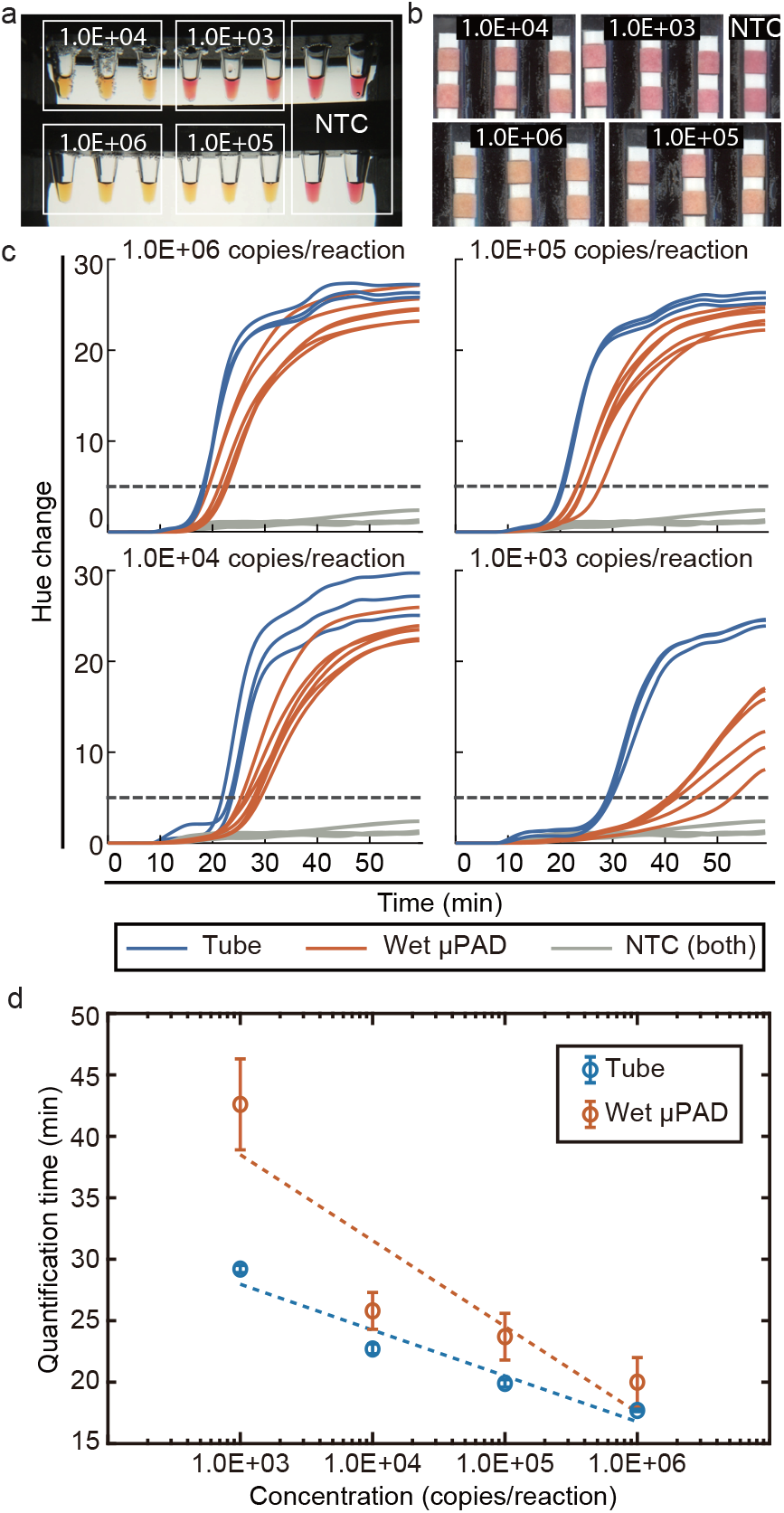
LAMP time differences between Tube and µPAD. (a) Experimental setup for Tube. (b) Experimental setup of µPAD. (c) LAMP curves of the reactions for each concentration. (d) Tq of Tube and µPAD with respect to concentration.

Figure 1b shows that the µPAD develops color more slowly. As shown in Figure 1b, even at 1.0E+06 copies, none of the six µPAD replicates show a fully converted hue after 30 min. Devices with lower template concentrations exhibit progressively weaker transitions. Both protocols are heated under identical conditions, revealing an intrinsic kinetic limitation in the µPAD reaction.

Hue-versus-time curves quantify the reaction rate and yield Tq values (Figure 1c). Tube reactions (blue) reach the quantification threshold (black dashed line) more rapidly than µPADs (vermil-ion). Figure 1d summarizes these Tq values: µPADs yield 20, 23.7, 25.8, and 42.6 min, whereas Tubes reach Tq at 17.7, 19.9, 22.7, and 29.2 min for each genomic load (1.0E+06, 1.0E+05, 1.0E+04, and 1.0E+03 copies/reaction, respectively). Thus, µPADs operate 13%, 19%, 14%, and 46% more slowly than Tube reactions. The kinetic delay becomes increasingly pronounced as the template concentration decreases.

### 3.2 Heat-Transfer Behavior

The LAMP reaction can be conceptually divided into two distinct stages. The first stage is the initiation step, which involves primer strand invasion into the target duplex. Kinetic analysis by Dangerfield *et al*. showed that strand invasion is the rate-limiting step and occurs over several minutes before exponential amplification begins ^29^. The appropriate temperature is needed for thermal breathing and strand invasion ^35^.

A numerical study evaluates heat transfer in a submerged µPAD and Tube. Figures 2b and 2c illustrate the simulation domains and model configurations. Conjugate heat transfer is modeled in COMSOL Multiphysics (v6.1). Heat transfer in solid and fluid regions follows the transient energy equation without a volumetric heat source,

**Fig. 2.**
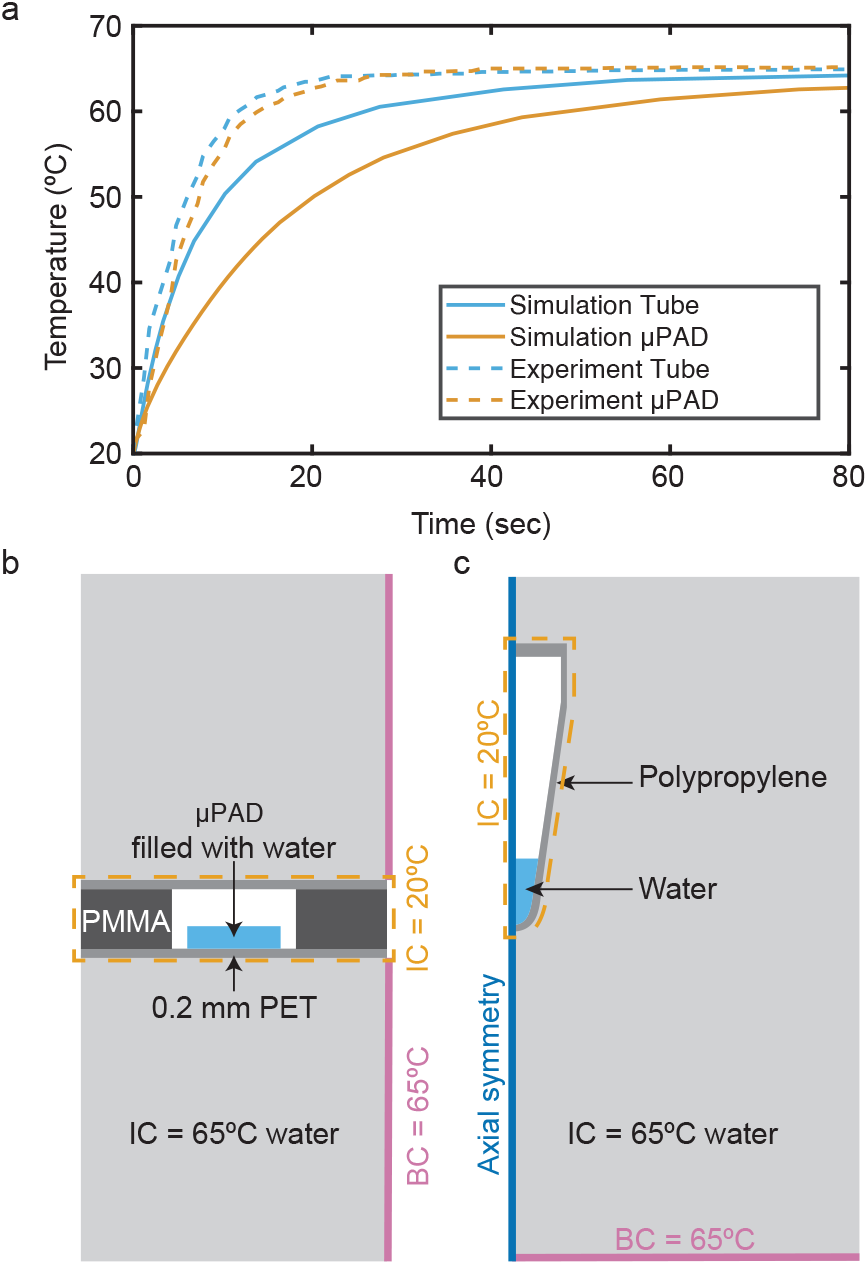
Experimental and numerical study of heat transfer. (a) Temperature with respect to time. (b) Simulation domain of the µPAD. (c) Simulation domain of the Tube

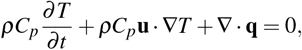

where *T* is temperature, *ρ* is density, *C*_*p*_ is specific heat capacity, and **u** is the flow velocity. Heat conduction follows **q** = −*k*_th_∇*T*, where *k*_th_ denotes thermal conductivity. Phase change is neglected. Laminar, incompressible Newtonian flow governs the fluid motion, and the heat-transfer and flow physics are fully coupled to account for convection. The computational domains consist of a two-dimensional µPAD and a two-dimensional axisymmetric tube.

As shown in Figure 2b, the surrounding medium consists of water initially at 65°C. The cartridge consists of polymethyl methacrylate (PMMA) with 0.2 mm polyethylene terephthalate (PET) adhesive layers on the top and bottom surfaces. A water-filled paper with 70% porosity lies on the lower PET layer, with the remaining enclosed region filled with air. The domain’s right boundary is fixed at 65°C, while all other boundaries are thermally insulated, −*n* **q** = 0. Figure 2c presents Tube geometry, consisting of a polypropylene tube containing 25 µL of water and the remaining volume filled with air. Tube initially remains at 20°C before being submerged in a 65°C bath. Axial symmetry is ap-plied to the left boundary, and the bottom boundary applies a 65°C heat source, with the remaining boundaries insulated.

Figure 2a compares the simulated and experimental temperatures of the µPAD and tube samples. The 25 µL Tube sample in the Tube reaches 60°C in under 26 s, whereas the µPAD reaches the same temperature at approximately 49 s. The difference between the two protocols (23 s) remains small relative to the experimentally observed reaction delay of up to 23 min between Tube LAMP and µPAD LAMP. Experimental results align with the simulations: tube reactions reach 60°C at 12 s, whereas µPADs reach it at approximately 14 s, as indicated by the dashed lines in Figure 2a. Consequently, the thermal lag between protocols, tens of seconds in simulation and only 2 seconds in measurement, is negligible compared to the 23 minute kinetic gap extracted from hue-based Tq analysis, indicating that post-warmup performance is unlikely to be governed by heat-up dynamics.

### 3.4 Effective Diffusion

The second stage of LAMP corresponds to the exponential amplification phase. In this regime, amplification proceeds through repeated primer binding to looped or hairpin structures, followed by strand-displacement synthesis. Savonnet *et al*. modeled this stage as being limited by primer hybridization, with an effective growth rate proportional to the primer concentration, *k*_obs_ = *k*_hyb_*C*_*P*_, where *k*_hyb_ is the second-order hybridization rate constant having upto 10 *µM*^−1^*s*^−1^ in solution and *C*_*P*_ is the primer concentration ^36^. As a result, *k*_obs_ in our case (primer concentration is 2 *µM*), resulting in *k*_obs_ ≈ 20 *s*^−1^

A qualitative assessment of the dominance of material transport over reactions is based on the Damköhler number (*Da*). Considering the reaction-diffusion equation,

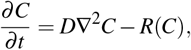

and assuming a first-order reaction *R* = *k*_obs_*C*, introduction of characteristic scales for concentration (*C*_0_) and length (*L*) yields

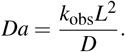

Using characteristic values, *D* ~ 10^−10^ m^2^ s^−1^ for primers ^37,38^, and *L* ~ 10 *µ*m ^39^, the resulting *Da* is on the order of 10. Thus, the exponential stage may operate near or above the transport-reaction transition *Da >* 1. In porous paper substrates, where the effective diffusion coefficient is reduced relative to free solution, *Da* is expected to increase further, potentially leading to diffusion-influenced amplification dynamics.

Diffusion in the µPAD depends on the porous microstructure. Effective diffusivity accounts for obstruction by fibers and is commonly described as

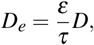

where *ε* is porosity and *τ* is tortuosity ^40^. Tortuosity is defined as *τ* = (⟨*L*_*d*_ ⟩*/L*_*s*_)^2^, where ⟨*L*_*d*_ ⟩ is the mean diffusive pathway length and *L*_*s*_ is the straight-line distance ^40^. Several empirical models estimate *τ* in saturated porous media, including the Weissberg model *τ* = 1 − 0.5 ln *ε* ^41^, the Archie model *τ* = *ε*^−*β* 42^, the Millington-Quirk model *τ* = *ε*^−1*/*3 43^, and the Delgado model *τ* = *ε*^−0.4 44^. Although these models differ in form, they consistently demonstrate reduced diffusivity relative to free solution. These relationships enable qualitative estimation of diffusion reduction in µPADs, even though tortuosity also depends on microstructural features beyond porosity.

According to the manufacturer’s information, the properties of the paper we used are shown in Table 3^45^. The *ε* value of the paper, 75%, can be calculated using the formula *ε* = 1 −(*w/θ*)*/ρ*_*cotton*_, where *w* is the basis weight, *θ* is the thickness, and *ρ*_*cotton*_ is the density of cotton fiber. The effective diffusion calculated using the Weissberg, Millington-Quirk, and Delgado models is roughly two-thirds of the free-solution diffusivity. However, these porosity-only models may overestimate *D*_*e*_ in fibrous papers, where additional effects such as pore constrictivity, disconnected porosity, partial saturation, and solute-fiber interactions can further reduce the effective transport rate. Therefore, *D*_*e*_ ≈ 0.67*D* should be interpreted as an upper-bound estimate in µPAD substrates.

**Table 3.** Properties of the Ahlstrom chromatography electrophoresis and blotting paper grade 222.

| Thickness ($\theta$) | Basis Weight ($w$) | Density of cotton fiber ($\rho_{cotton}$) |
| --- | --- | --- |
| 0.81 mm | 0.302 kg/m<sup>2</sup> | 1500 kg/m<sup>3</sup> |

The practical implication is that tubes behave as well-mixed systems with short transport distances (*Da <* 1). In contrast, µPADs operate within a porous network that slows molecular motion and increases the effective encounter time for polymerase, primers, and template. In such environments, reduced effective diffusivity raises the Damköhler number above unity during the exponential stage, thereby causing the reaction to enter a diffusion-influenced regime in which spatial concentration gradients can form. Because our measured thermal transients are far too small to account for the minute-scale kinetic disparity, these transport considerations support the conclusion that restricted diffusion, rather than heating limitations, plays a substantial role in the delayed amplification observed in µPADs, particularly during early amplification near the detection limit.

### 3.4 Nonspecific Binding of Reactants on Fibers in µPAD

Reducing nonspecific binding of LAMP reagents to cellulose fibers provides another means to improve amplification speed on paper. Cellulose can adsorb DNA, polymerases, and other components, lowering their effective concentrations and slowing reaction initiation. Minimizing nonspecific binding, therefore, enhances both sensitivity and reaction kinetics ^46^. Blockers such as BSA are commonly used to occupy adsorption sites before amplification begins, while surfactants like Tween-20 reduce hydrophobic interactions ^47^, and trehalose helps stabilize biomolecules on dried substrates ^48^. Previous µPAD studies incorporate these additives to mitigate nonspecific binding and maintain reagent activity ^2^.

However, the effect of applying reagents to the paper before drying versus immediately before amplification has not been systematically evaluated. To address this, we compared two preparation schemes: “Wet µPAD” and “Dry µPAD.” In the Wet condition, all reagents and template DNA are deposited simultaneously onto the paper and immediately heated. In the Dry condition, all reagents except the template are applied first and allowed to dry for 24 hours at room temperature; template DNA is then added immediately before heating. Each condition includes six replicates per concentration, and all devices are enclosed in a single cartridge and incubated in a 65°C water bath. The NTC consists of four replicates, evenly divided between the two conditions.

Figure 3b displays representative colorimetric progressions at 15 min intervals for 1.0E+03 copies per reaction. The Wet µPAD exhibits a slower hue shift than the Dry µPAD, with clear differences emerging between 45 and 60 min. Hue-versus-time plots in Figure 3c quantify these observations. For 1.0E+06 copies/reaction (top-left panel), vermilion traces correspond to Wet µPADs and light-blue traces to Dry µPADs, while gray curves (NTC) remain flat. Remaining panels show analogous data for 1.0E+05 through 1.0E+03 copies/reaction.

**Fig. 3.**
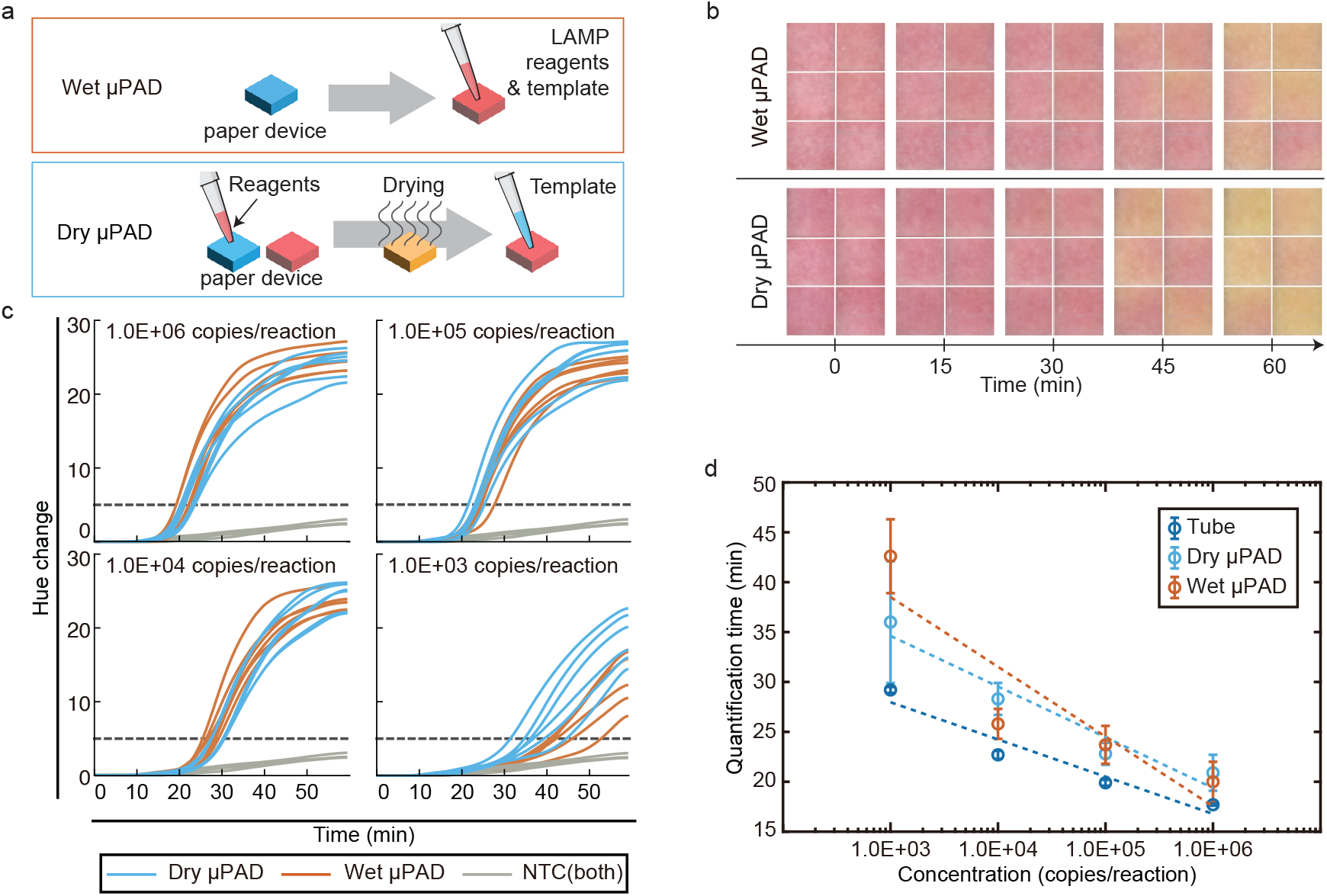
Experimental setup and result of the experiment with Wet µPAD and Dry µPAD. (a) Preparation methods for controls. The top shows the preparation procedure for Wet µPAD, while the bottom shows Dry µPAD. (b) Scanned images of paper devices with different protocols over time. The concentration is 1.0E+03 copies/reaction. (c) LAMP curves for replicates with respect to time. (d) Tq with respect to the concentration of template copies/reaction.

The Tq values (Figure 3d) indicate similar performance between Wet and Dry µPADs for 1.0E+06, 1.0E+05, and 1.0E+04 copies. Tq of Wet µPADs are 20, 23.7, and 25.8 min, respectively. Tq of Dry µPADs are 20.9, 22.8, and 28.3 min, respectively. At 1.0E+03 copies, however, the Wet µPAD shows a pronounced delay, with a Tq of 42.6 min compared to 36 min for the Dry condition. At low target concentrations, adsorption removes a disproportionate fraction of LAMP reactants, so even modest nonspecific surface binding can significantly impair amplification.

These findings demonstrate that the order and stage of reagent addition influence amplification kinetics in the µPAD device. Among the blocking components, BSA is particularly sensitive to when it is applied: pre-coating the fiber surface more effectively suppresses nonspecific binding. In contrast, Tween-20 primarily modulates hydrophobic interactions, and trehalose stabilizes reagents during drying; varying the stage at which they are added does not produce a measurable effect compared with BSA pre-coating.

We next test whether applying BSA before drying alters amplification behavior. Figure 4a outlines two different control preparations. In the “Wet BSA” condition, all LAMP reagents except BSA and template are first deposited and dried for 24 hours; BSA and template are then added immediately before heating. In the “Dry BSA” condition, BSA is first deposited and dried for 24 hours, and the remaining reagents and template are added afterwards. This design isolates whether pre-saturating cellulose fibers with BSA enhances blocking efficiency at the onset of amplification.

**Fig. 4.**
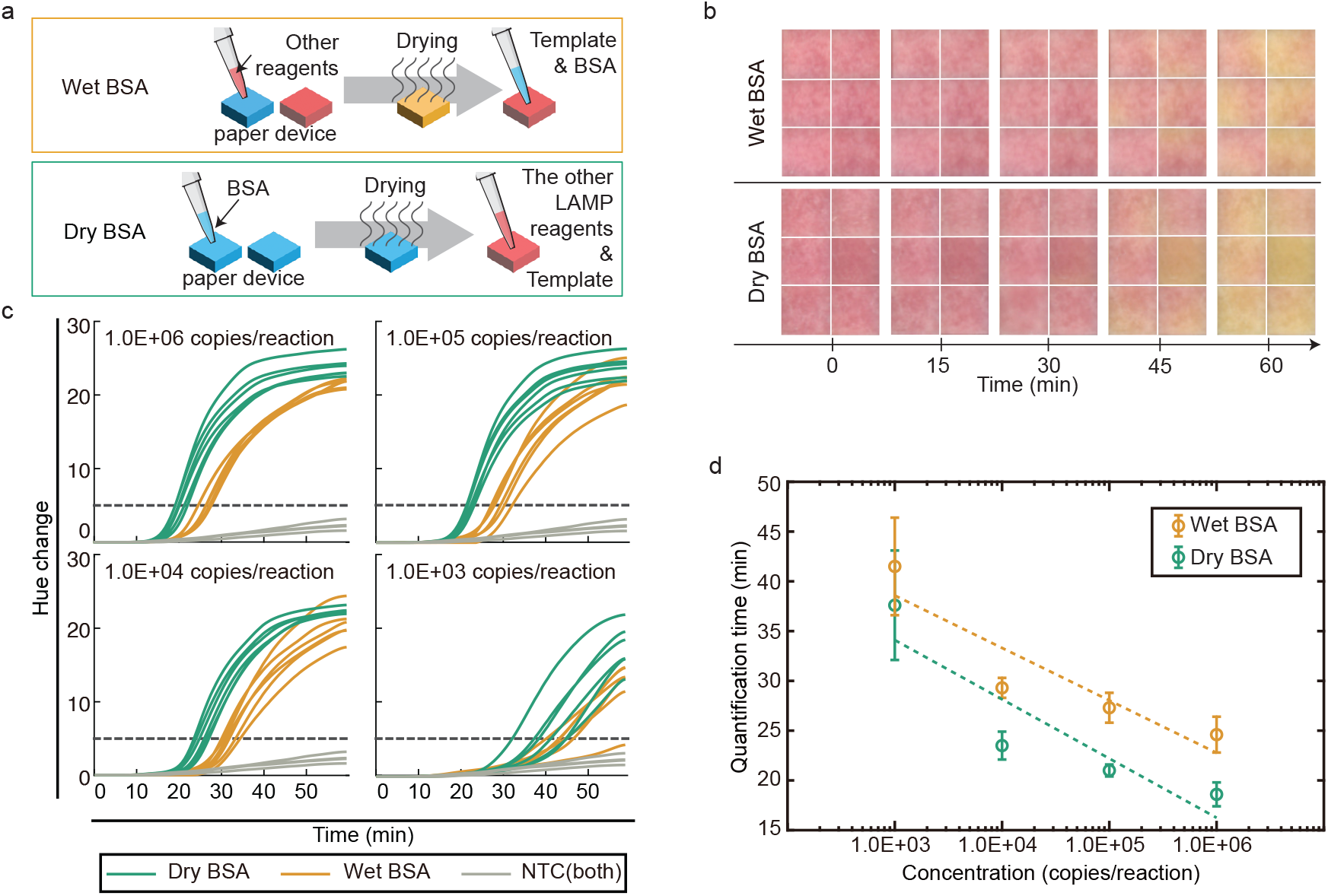
Experimental setup and result of the experiment with Wet BSA and Dry BSA. (a) Preparation methods for controls. The top panel shows the preparation procedure for Wet BSA, while the bottom panel shows that for Dry BSA. (b) The scanned images of paper devices with different protocols over time. The concentration is 1.0E+03 copies/reaction. (c) LAMP curves for replicates with respect to time. (d) Tq with respect to the concentration of template copies/reaction.

The colorimetric images in Figure 4b show faster hue transitions for Dry BSA compared to Wet BSA, particularly at 1.0E+03 copies per reaction. The hue-versus-time curves in Figure 4c reinforce these visual differences: green traces (Dry BSA) cross the threshold earlier than orange traces (Wet BSA), while gray traces (NTC) are flat. Tq analysis in Figure 4d summarizes performance across concentrations. Both conditions display the expected inverse dependence of Tq on template concentration. However, Dry BSA consistently yields shorter Tq values of 18.6, 21, 23.5, and 37.6 min at concentrations ranging from 1.0E+6 to 1.0E+3, respectively. In comparison, Wet BSA shows Tq of 24.6, 27.3, 29.3, and 41.5 min, respectively. Dry BSA reduces Tq 10-30% relative to Wet BSA.

## 4 Discussion

The results clarify why LAMP amplification on paper lags behind liquid reactions. The first stage of the LAMP is the initiation step, which depends on dsDNA thermal breathing and temperature-dependent strand invasion. ^29,35^. Although approaching 65°C promotes the initiation step, both numerical and experimental measurements show that µPADs and tubes warm within seconds, indicating that bulk heating cannot explain the multi-minute kinetic gap ^31^. Instead, reduced effective diffusion and nonspecific adsorption dominate. Using pore-scale length scales, macromolecular diffusivities, and a reaction rate, the Damköhler number reaches ~10, indicating diffusion-influenced kinetics in µPAD ^36^. For Ahlstrom grade 222 (porosity *ε* ≈ 0.75), empirical tortuosity relations ^41,43,44^ predict maximum diffusion of *D*_*e*_ ≈ (2*/*3)*D*, consistent with the observed transport penalty.

By contrast, Tube operates under well-mixed, reaction-limited conditions, indicating that diffusion or material transport is faster than the chemical reaction rate (*Da <* 1) in Tube. If the reaction were still under the diffusion-limited region in Tube, the Tq value of Tube should be less than 24 min, which is 2/3 of Tq at µPAD, 36 min. The results from the Tube reaction (29.2 min) indicate a lag due to the chemical reaction itself.

Surface interactions add another constraint. Cellulose fibers adsorb nucleic acids and proteins ^34^. When BSA is applied before the other reagents (Dry BSA), Tq consistently shifts earlier compared to Wet conditions, aligning with prior reports on the value of blocking agents in µPAD amplification ^46,49^. Pre-coating saturates adsorption sites, reducing losses that otherwise penalize reactions.

Among the tested preparation protocols, “Dry BSA”, which provides the most effective blocking of nonspecific adsorption, produces the lowest Tq values in the high-copy regime (1E+04 to 1E+06 copies/reaction). This result indicates that, at high template concentrations, diffusion is no longer limiting, and nonspecific binding becomes the dominant source of delay, whereas pre-coating the substrate with BSA effectively mitigates this penalty. Consistently, Dry BSA shows only a 4.7% Tq delay relative to Tube in the high-copy regime. In comparison, the “Dry µPAD” condition, where BSA is applied together with the other LAMP reagents, exhibits a 19.1% delay. The least effective condition, “Wet BSA”, in which BSA is added after the other reactants, shows a 35.1% delay. In low-copy regimes (1E+03 copies/reaction), diffusion masks the benefits of blocking. It is evidenced by the increased lag of “Dry BSA” vs. Tube reaction, from 4.7% to 28.8%, from high copies/reaction to low copies/reaction, respectively.

Although heat transfer, transport, and surface phenomena can couple, their time scales differ substantially. Heating equilibrates in seconds, whereas diffusion and adsorption collectively delay amplification by minutes. Thus, interventions targeting mass transport and adsorption, not further thermal optimization, are most impactful. These insights suggest practical design rules for µPAD LAMP. Substrates with higher porosity and lower tortuosity may alleviate diffusion limitations. BSA pre-coating should be implemented routinely before adding template and polymerase. Device geometries that shorten diffusion paths may also enhance performance without sacrificing operational simplicity.

Limitations include the use of empirical tortuosity relations that depend on fiber morphology and saturation, as well as variability introduced by drying or reagent distribution. Future work should experimentally quantify adsorption isotherms, measure *D*_*e*_, and develop coupled reaction-diffusion-adsorption models validated against hue-time trajectories. These efforts will refine substrate engineering and blocking strategies that further close the gap between µPAD and Tube LAMP.

## Conclusions

This study demonstrates that the primary barriers to rapid µPAD LAMP amplification arise not from heat transfer limitations but from mass transport and surface interactions inherent to porous cellulose substrates. Once thermal equilibrium is reached, amplification on paper is slowed by restricted diffusion and by nonspecific adsorption of primers, polymerase, and template to cellulose fibers. By experimentally isolating these contributions, we show that substrate porosity, tortuosity, and adsorption behavior collectively govern the kinetic lag relative to well-mixed tube reactions. The balance between diffusion limitation and surface adsorption depends on the input copy number. At high template concentrations, diffusion ceases to be rate-limiting, and nonspecific adsorption dominates the residual delay. Pre-coating with BSA mitigates early adsorption losses, reducing the Tq difference to 5% relative to Tube reactions. At low copy numbers, diffusion constraints reassert themselves and outweigh the benefit of blocking, leading to substantial delays even under the Dry BSA protocol.

These findings provide clear design principles for faster and more consistent µPAD LAMP assays: selecting substrates with lower tortuosity, integrating routine pre-coating strategies, and minimizing diffusion path lengths are critical for performance gains. More broadly, these results underscore the importance of coupling material selection with surface conditioning to overcome the intrinsic transport and adsorption penalties of fibrous matrices. Such strategies will be essential for advancing the reliability and speed of next-generation point-of-need nucleic acid diagnostics on paper.

## Data availability

All data generated or analyzed during this study are included in this published article. Analysis scripts are available from the corresponding author upon reasonable request.

## Acknowledgements

Funding for this project was provided by the American Rescue Plan Act through USDA APHIS. The findings and conclusions in this publication are those of the author(s) and should not be construed to represent any official USDA or U.S. Government determination or policy. We sincerely thank Dr. Jiangshan Wang for his valuable advice on experimental planning. We also gratefully acknowledge Cindy Peres for her training in general nucleic acid amplification techniques.

## References

1 T. Kang, J. Lu, T. Yu, Y. Long and G. Liu, Advances in nucleic acid amplification techniques (NAATs): COVID-19 point-of-care diagnostics as an example, 2022.

2 J. L. Davidson, J. Wang, M. K. Maruthamuthu, A. Dextre, A. Pascual-Garrigos, S. Mohan, S. V. S. Putikam, F. O. I. Osman, D. McChesney, J. Seville and M. S. Verma, Biosensors and Bioelectronics: X, 2021, 9, 100076.

3 J. Wang, A. Dextre, A. Pascual-Garrigos, J. Levi Davidson, M. K. Maruthamuthu, D. McChesney, J. Seville and M. S. Verma, MethodsX, 2021, 8, 101586.

4 T. Notomi, H. Okayama, H. Masubuchi, T. Yonekawa, K. Watanabe, N. Amino and T. Hase, Nucleic Acids Research, 2000, 28, e63–e63.

5 K. Nagamine, T. Hase and T. Notomi, Molecular and Cellular Probes, 2002, 16, 223–229.

6 Y. Zhao, F. Chen, Q. Li, L. Wang and C. Fan, Chemical Reviews, 2015, 115, 12491–12545.

7 K. J. Moore, J. Cahill, G. Aidelberg, R. Aronoff, A. Bektaş, D. Bezdan, D. J. Butler, S. V. Chittur, M. Codyre, F. Federici, N. A. Tanner, S. W. Tighe, R. True, S. B. Ware, A. L. Wyllie, E. E. Afshin, A. Bendesky, C. B. Chang, R. I. Rosa, E. Elhaik, D. Erickson, A. S. Goldsborough, G. Grills, K. Hadasch, A. Hayden, S. Y. Her, J. A. Karl, C. H. Kim, A. J. Kriegel, T. Kunstman, Z. Landau, K. Land, B. W. Langhorst, A. B. Lindner, B. E. Mayer, L. A. McLaughlin, M. T. McLaughlin, J. Molloy, C. Mozsary, J. L. Nadler, M. D’silva, D. Ng, D. H. O’connor, J. E. Ongerth, O. Osuolale, A. Pinharanda, D. Plenker, R. Ranjan, M. Rosbash, A. Rotem, J. Segarra, S. Schürer, S. Sherrill-Mix, H. Solo-Gabriele, S. To, M. C. Vogt, A. D. Yu and C. E. Mason, Journal of Biomolecular Techniques, 2021, 32, 228–275.

8 A. Pascual-Garrigos, M. K. Maruthamuthu, A. Ault, J. L. Davidson, G. Rudakov, D. Pillai, J. Koziol, J. P. Schoonmaker, T. Johnson and M. S. Verma, Veterinary Research, 2021, 52, 126.

9 J. Wang, J. Davidson, S. Kaur, A. Dextre, M. Ranjbaran, M. Kamel, S. Athalye and M. Verma, Biosensors, 2022, 12, 1094.

10 J. Wang, M. Ranjbaran, A. Ault and M. S. Verma, Food Microbiology, 2023, 110, 104173.

11 J. Wang, S. Kaur, A. Kayabasi, M. Ranjbaran, I. Rath, I. Benschikovski, B. Raut, K. Ra, N. Rafiq and M. S. Verma, Biosensors and Bioelectronics, 2024, 259, 116374.

12 M. Kamel, J. L. Davidson, J. M. Schober, G. S. Fraley and M. S. Verma, Scientific Reports, 2025, 15, 12110.

13 B. Ahmed, B. Raut, A. Pauley, J. L. Davidson, S. Yang and M. S. Verma, Biosensors and Bioelectronics, 2025, 287, 117690.

14 B. Raut, G. Palla, V. Kumar, A. Fleck, B. Ahmed, J. L. Davidson, R. F. Relich, J. P. Schoonmaker, J. A. Pasternak and M. S. Verma, bioRxiv, 2026, 2026.01.09.696240.

15 B. Raut, G. Palla, J. Wang, S. Campbell, V. Kumar, K. Ra and M. S. Verma, bioRxiv, 2026, 2026.01.15.699357.

16 H. Q. Nguyen, V. D. Nguyen, H. Van Nguyen and T. S. Seo, Scientific Reports 2020 10:1, 2020, 10, 15123–.

17 G. Papadakis, A. K. Pantazis, N. Fikas, S. Chatziioannidou, V. Tsiakalou, K. Michaelidou, V. Pogka, M. Megariti, M. Vardaki, K. Giarentis, J. Heaney, E. Nastouli, T. Karamitros, A. Mentis, A. Zafiropoulos, G. Sourvinos, S. Agelaki and E. Gizeli, Scientific Reports 2022 12:1, 2022, 12, 3775–.

18 J. C. Linnes, A. Fan, N. M. Rodriguez, B. Lemieux, H. Kong and C. M. Klapperich, RSC Advances, 2014, 4, 42245–42251.

19 D. Das, C. W. Lin and H. S. Chuang, Biosensors, 2022, 12, 1068.

20 K. Kaarj, P. Akarapipad and J. Y. Yoon, Scientific Reports, 2018, 8, 12438.

21 B. S. Batule, Y. Seok and M. G. Kim, Biosensors and Bioelectronics, 2020, 151, 111998.

22 S. Roy, N. F. Mohd-Naim, M. Safavieh and M. U. Ahmed, ACS Sensors, 2017, 2, 1713–1720.

23 B. Pang, K. Fu, Y. Liu, X. Ding, J. Hu, W. Wu, K. Xu, X. Song, J. Wang, Y. Mu, C. Zhao and J. Li, Analytica Chimica Acta, 2018, 1040, 81–89.

24 T. N. D. Trinh and N. Y. Lee, Lab on a Chip, 2019, 19, 1397– 1405.

25 K. T. L. Trinh, T. N. D. Trinh and N. Y. Lee, Biosensors and Bioelectronics, 2019, 135, 120–128.

26 M. Zhang, J. Liu, Z. Shen, Y. Liu, Y. Song, Y. Liang, Z. Li, L. Nie, Y. Fang and Y. Zhao, BMC Microbiology, 2021, 21, 197.

27 X. Ye, J. Xu, L. Lu, X. Li, X. Fang and J. Kong, Analytica Chimica Acta, 2018, 1018, 78–85.

28 N. Sritong, M. Sala de Medeiros, L. A. Basing and J. C. Linnes, Lab on a Chip, 2023, 23, 888–912.

29 T. L. Dangerfield, I. Paik, S. Bhadra, K. A. Johnson and A. D. Ellington, Nucleic Acids Research, 2023, 51, 488–499.

30 D. Das and S. Namboodiri, Chemical Engineering Science, 2021, 229, 116130.

31 D. Das, M. Masetty and A. Priye, Chemosensors 2023, Vol. 11,, 2023, 11, 163.

32 I. A. Badruddin, Azeem, T. M. Yunus Khan and M. A. Ali Baig, Materials Today: Proceedings, 2020, 24, 1318–1321.

33 D. Domiri Ganji and E. Mohseni Languri, Heat Transfer - Mathematical Modelling, Numerical Methods and Information Technology, IntechOpen, London, 2011, ch. 27, pp. 631–642.

34 W. Norde, Colloids and Surfaces B: Biointerfaces, 2008, 61, 1– 9.

35 J. H. Jeon, W. Sung and F. H. Ree, Journal of Chemical Physics, 2006, 124, 164905.

36 M. Savonnet, M. Aubret, P. Laurent, Y. Roupioz, M. Cubizolles and A. Buhot, Biosensors, 2022, 12, 346.

37 S. Kondrat and M. N. Popescu, Physical Chemistry Chemical Physics, 2019, 21, 18811–18815.

38 G. L. Lukacs, P. Haggie, O. Seksek, D. Lechardeur, N. Freedman and A. S. Verkman, Journal of Biological Chemistry, 2000, 275, 1625–1629.

39 H. Aslannejad and S. M. Hassanizadeh, Transport in Porous Media 2017 120:1, 2017, 120, 67–81.

40 B. Ghanbarian, A. G. Hunt, R. P. Ewing and M. Sahimi, Soil Science Society of America Journal, 2013, 77, 1461–1477.

41 H. L. Weissberg, Journal of Applied Physics, 1963, 34, 2636– 2639.

42 G. E. Archie, SPE reprint series, 2003, 9–18.

43 R. J. Millington and J. P. Quirk, Trans. Faraday Soc., 1961, 57, 1200–1207.

44 J. M. Delgado, Canadian Journal of Chemical Engineering, 2006, 84, 651–655.

45 Ahlstrom, Chromatography and Blotting Transfers, high quality cotton paper, 2023, https://ahlstrom-munksjo.showpad.com/share/mK0HxCgLgOU98L0q7n9cs.

46 Y. Liu, L. Zhan, Z. Qin, J. Sackrison and J. C. Bischof, ACS Nano, 2021, 15, 3593–3611.

47 K. Kluge, S. J. Squillace, S. J. Mao and B. A. Kottke, Biochimica et Biophysica Acta (BBA)/Lipids and Lipid Metabolism, 1980, 620, 447–453.

48 A. Dubrow, B. Zuniga, E. Topo and J. H. Cho, ACS Omega, 2022, 7, 9206–9211.

49 A. R. von Stockert, A. Luongo, M. Langhans, T. Brandstetter, J. Rühe, T. Meckel and M. Biesalski, Sensors, 2021, 21, 6348.

